# Biologically-informed deep neural networks provide quantitative assessment of intratumoral heterogeneity in post-treatment glioblastoma

**DOI:** 10.1101/2022.12.20.521086

**Authors:** Hairong Wang, Michael G Argenziano, Hyunsoo Yoon, Deborah Boyett, Akshay Save, Petros Petridis, William Savage, Pamela Jackson, Andrea Hawkins-Daarud, Nhan Tran, Leland Hu, Osama Al Dalahmah, JeffreyN. Bruce, Jack Grinband, Kristin R Swanson, Peter Canoll, Jing Li

## Abstract

Intratumoral heterogeneity poses a significant challenge to the diagnosis and treatment of glioblastoma (GBM). This heterogeneity is further exacerbated during GBM recurrence, as treatment-induced reactive changes produce additional intratumoral heterogeneity that is ambiguous to differentiate on clinical imaging. There is an urgent need to develop non-invasive approaches to map the heterogeneous landscape of histopathological alterations throughout the entire lesion for each patient. We propose to predictively fuse Magnetic Resonance Imaging (MRI) with the underlying intratumoral heterogeneity in recurrent GBM using machine learning (ML) by leveraging image-localized biopsies with their associated locoregional MRI features. To this end, we develop BioNet, a biologically-informed neural network model, to predict regional distributions of three tissue-specific gene modules: proliferating tumor, reactive/inflammatory cells, and infiltrated brain tissue. BioNet offers valuable insights into the integration of multiple implicit and qualitative biological domain knowledge, which are challenging to describe in mathematical formulations. BioNet performs significantly better than a range of existing methods on cross-validation and blind test datasets. Voxel-level prediction maps of the gene modules by BioNet help reveal intratumoral heterogeneity, which can improve surgical targeting of confirmatory biopsies and evaluation of neuro-oncological treatment effectiveness. The non-invasive nature of the approach can potentially facilitate regular monitoring of the gene modules over time, and making timely therapeutic adjustment. These results also highlight the emerging role of ML in precision medicine.

## Introduction

Glioblastoma (GBM) exhibits pronounced intratumoral heterogeneity, which can confound diagnosis and clinical management, and is a leading driver of tumor recurrence (1, 2). Treatment-induced reactive changes further exacerbate intratumoral heterogeneity (3, 4). Because histopathological and molecular analyses are limited by sparse biopsy sampling, there is a significant need to develop non-invasive approaches to map the heterogeneous landscape of histopathological alterations throughout the entire lesion. Such advancements would improve surgical targeting of confirmatory biopsies and non-invasive assessment of neuro-oncological treatment response, thereby informing subsequent therapeutic strategies. Radiogenomics is a growing research field, which seeks to develop machine learning (ML) models to predict cellular, molecular and genetic characteristics of tumors based on Magnetic Resonance Imaging (MRI) and other imaging types (2, 5, 6). Radio(gen)omics methods have been shown to accurately predict not only diversity in tumor cell density associated with diffuse invasion into the brain parenchyma peripheral to the frank lesion seen on MRI (7–9), but also abnormalities in hallmark genes such as EGFR, PDGFRA, and PTEN (10–13), IDH mutation status (14–18), and MGMT methylation status based on radiographic features (13–15). These studies represent examples in which the prediction provides a single categorical label per tumor using imaging features that span the entire lesion.

However, precise representations of intratumoral heterogeneity require *voxel-wise* labels (e.g., image-localized biopsies) that reflect local or regional characteristics of the lesion. A major challenge for such prediction is the lack of large image-localized biopsy datasets (19, 20) to train deep learning (DL) models that are well-known to be heavily-parameterized and data-hungry. Creation of large training datasets is limited by various factors such as the invasiveness and high expense of sample acquisition, need of highly-specialized experts to create accurate labels, and difficulty in patient recruitment (19). Moreover, the lack of large datasets has severely limited the number of studies focusing on predicting regional characteristics within each lesion, which are crucial for revealing intratumoral heterogeneity. A few studies have developed MRI-biology fusion models to predict regional cell density (21–26) or regional copy number variation of individual driver genes such as EGFR, PDGFRA, and PTEN, in untreated, primary GBM (27, 28). In recurrent GBM (recGBM), however, treatment-induced reactive changes lead to additional intratumoral heterogeneity and the related additional complexity in tissue composition makes prediction of gene modules more difficult (29).

In this study, we compiled a unique dataset that included multi-region biopsy samples and MRI from recGBM patients. The dataset consisted of derived measurements for three gene modules, from each biopsy, by combining data of individual gene expressions from RNA sequencing and cellular composition patterns from immunohistochemistry (IHC). The three gene modules identified through gene ontology analysis include: proliferative (**Pro**), associated with proliferation and cell cycle ontologies indicative of recurrent tumor; inflammatory (**Inf**), linked to cytokine production and immune response, representative of treatment-induced reactive cells; and neuronal (**Neu**), related to neuronal signaling, reflecting infiltrated brain tissue. Assessing the gene modules of GBM has significant clinical value and has drawn much attention recently (30). For recGBM, the ability to differentiate proliferative/recurrent tumor and treatment-induced reactive/inflammatory cells (two primary gene modules in our dataset) is crucial for evaluating treatment effectiveness. However, such differentiation is notoriously difficult in clinical practice due to their indistinguishable appearances on MRI. Even among seasoned practitioners, accurately distinguishing between proliferative/recurrent tumors and treatment-induced reactive/inflammatory cells remains an elusive task. Currently, the sole method for distinguishing between these two gene modules is obtaining biopsies and conducting comprehensive transcriptomic and immunohistochemical profiling. However, biopsies, the gold-standard approach, can only cover a few sparse regions, leaving substantial regions within the lesion unexamined and the differentiation in these regions is nearly equivalent to a random guess. Therefore, our unique dataset, comprising a development cohort and a test cohort, facilitated the first-ever development of a non-invasive approach based on MRI and DL to predict voxel-level gene modules throughout the entire lesion for each patient.

To tackle the inherent challenge of limited training data from biopsy samples, we proposed BioNet, a novel unified framework whose learning capacity is significantly augmented by integrating multiple implicit and qualitative biological domain knowledge. The integration of biological/biomedical domain knowledge, such as biological principles, empirical models, simulations, and knowledge graphs, can provide a rich source of information (pseudo data) to help alleviate the data shortage in training DL models. Various approaches have been proposed for integrating domain knowledge, depending on its form. For example, some researchers proposed to use the knowledge of biological pathways to guide the design of DL architecture (31, 32). In certain biomedical domains, knowledge exists in the form of algebraic equations that capture biological principles, which were integrated with DL architecture or loss functions (23, 33). Some researchers proposed to integrate knowledge about feature behavior as attribution priors into DL training (34). However, existing methods lack the ability to simultaneously incorporate multiple implicit, qualitative domain knowledge that is difficult to describe in mathematical formulations, making them unsuitable for our problem (35). To fully harness the potential of this type of domain knowledge, BioNet integrated several strategies. Firstly, it creates large virtual biopsy datasets based on domain knowledge to pre-train the DL model, enabling it to learn generalizable feature representations that can be transferred to the downstream task based on real biopsy samples. Secondly, BioNet adopts a hierarchical design inspired by domain knowledge, considering the interaction between gene modules and their conditional relationships. Lastly, BioNet employs a knowledge attention loss function that combines data-driven and knowledge-driven components, penalizing violations of domain knowledge on unlabeled samples. These strategies collectively empower BioNet to effectively leverage and incorporate domain knowledge into the learning process.

In summary, by leveraging a uniquely compiled dataset, this study is the first of its kind that developed a non-invasive approach for quantifying intratumoral heterogeneity in the recGBM setting. The focus was to predict the regional distributions of gene modules representing proliferative/recurrent tumor and treatment-induced reactive/inflammatory cells using MRI. The mapping of these gene module distributions throughout the entire lesion for each patient offers valuable clinical benefits. It can assist in the identification of locations within a lesion for confirmatory biopsy sampling. With BioNet’s guidance, the likelihood of sampling locations with the desired gene modules is significantly improved, offering a more reliable alternative to the current sampling approach. It can also assist clinicians in evaluation of treatment effectiveness. The non-invasive nature of the approach can potentially facilitate regular monitoring of the gene modules over time, enabling identification of treatment response or resistance and making timely therapeutic adjustment. By gaining granularity from regional assessment, clinicians may gain better understanding of patient-specific nuances and tailor treatment more individually.

## Results

This study involved a developmental cohort and a test cohort, referred to as cohort A and B hereafter. **Fig. 1** presents an overview of the application of BioNet in assisting the assessment of treatment responses and informing subsequent therapy decisions.

**Fig. 1.**
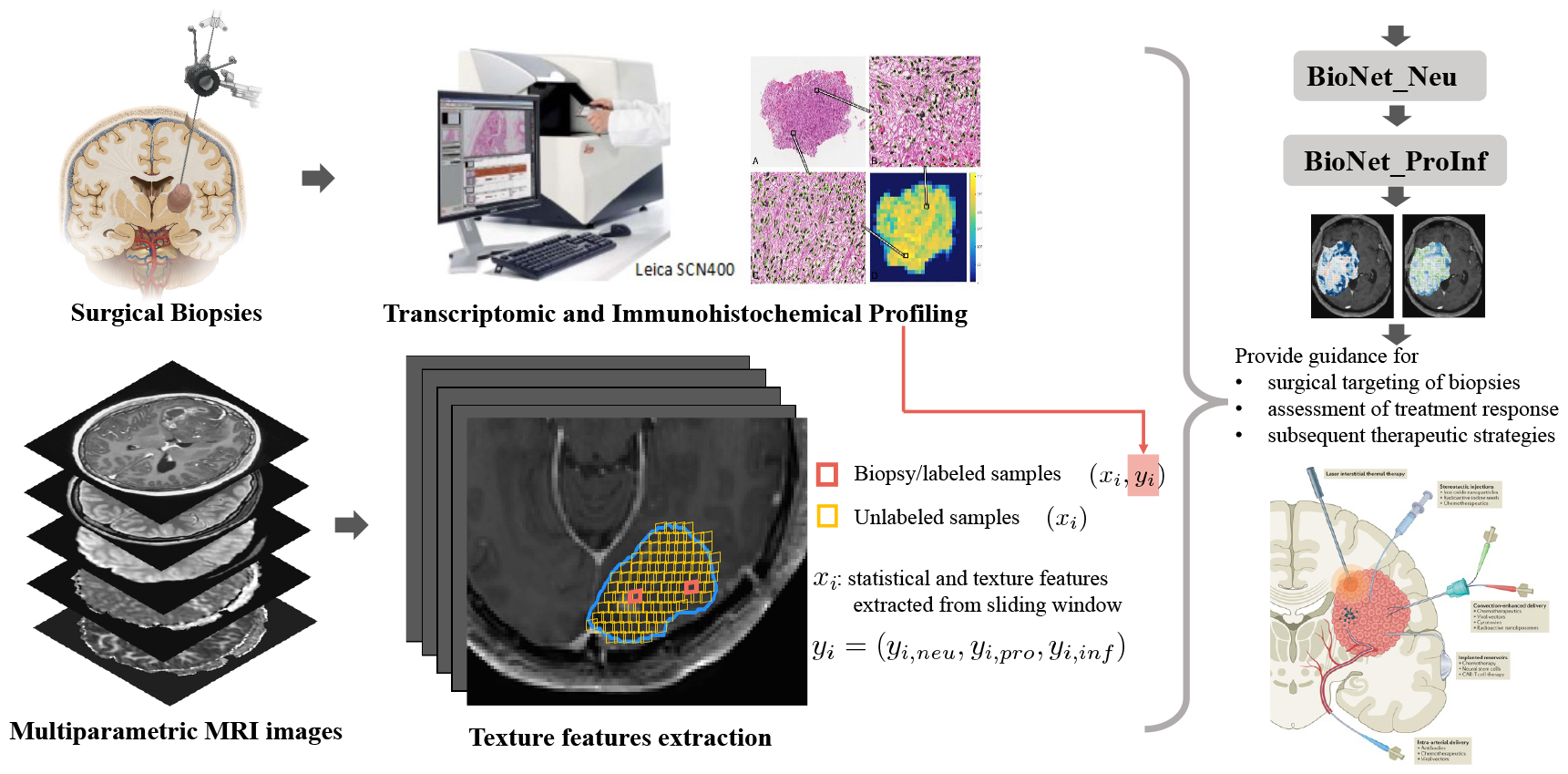
Overview of the application of BioNet in assisting the assessment of treatment responses and informing subsequent therapy decisions. Our datasets comprise two types of data: sparsely labeled data from biopsy locations, and abundant unlabeled data from all locations throughout the entire brain. The labels for biopsy samples (*y*_*i*_) are obtained through comprehensive transcriptomic and immunohistochemical profiling. The input features (*x*_*i*_) for all samples, both labeled and unlabeled, which are utilized in the training and testing of BioNet, are extracted from multiparametric MRI. Within the tumoral Area of Interest (AOI) of each patient (blue outline), local regions (small squares) were created by sliding windows according to the physical size of surgical biopsies. Statistical and texture features (*x*_*i*_) were computed based on multiparametric MRI within the sliding windows at biopsy locations (red, a few)and remaining unlabeled locations (yellow, abundant). Labeled samples, along with selectively chosen unlabeled samples, are employed in the training of BioNet. Once adequately trained, BioNet is capable of generating *voxel-level* prediction maps for proliferative/recurrent tumors (Pro) and treatment-induced reactive/inflammatory cells (Inf) respectively within AOI. These prediction maps yield crucial insights into the gene status at the voxel level throughout the entire tumor. The picture of Surgical Biopsy is adapted from (36). The picture at the right bottom about local therapies for GBM is adapted from (37).

### Biologically-informed design principles for BioNet

Our goal is to develop a model, which can accurately predict Pro (*y*_*i,pro*_) and Inf (*y*_*i,inf*_) scores for each region *i* within a tumoral Area of Interest (AOI) based on regional MRI features (*x*_*i*_), for individual patients. Obtaining regional distributions of these scores throughout the AOI can help improve surgical targeting of confirmatory biopsies and evaluate treatment effectiveness for each patient, thereby informing subsequent therapeutic strategies. Training an effective predictive model solely on labeled/biopsy samples presents inherent limitations due to the extremely small sample size. Conversely, there exists a substantial volume of unlabeled samples from AOI that could be strategically utilized. Thus, by leveraging unlabeled samples, BioNet is proposed to integrate domain knowledge through unique framework design.

Specifically, our domain knowledge, based on fundamental understanding of the neuropathological relationships between the three gene modules (30), reveals two key relationships between Pro and Inf conditional on the status of Neu (**Fig. 2a**): (1) We know that genes in the Neu module are enriched in normal brain tissue and depleted in lesional brain tissue. Thus, samples with high Neu tend to have low Pro and low Inf. (2) We also know that in samples with low Neu, the lesional component of the tissue comprises a mixture of proliferative tumor and inflammatory response. Thus, if a sample has more proliferative tumor, i.e., Pro high, it is likely to have less inflammatory response, i.e., Inf Low, and vice versa. This implies that samples with low Neu values are inclined to exhibit a negative correlation between Pro and Inf.

**Fig. 2.**
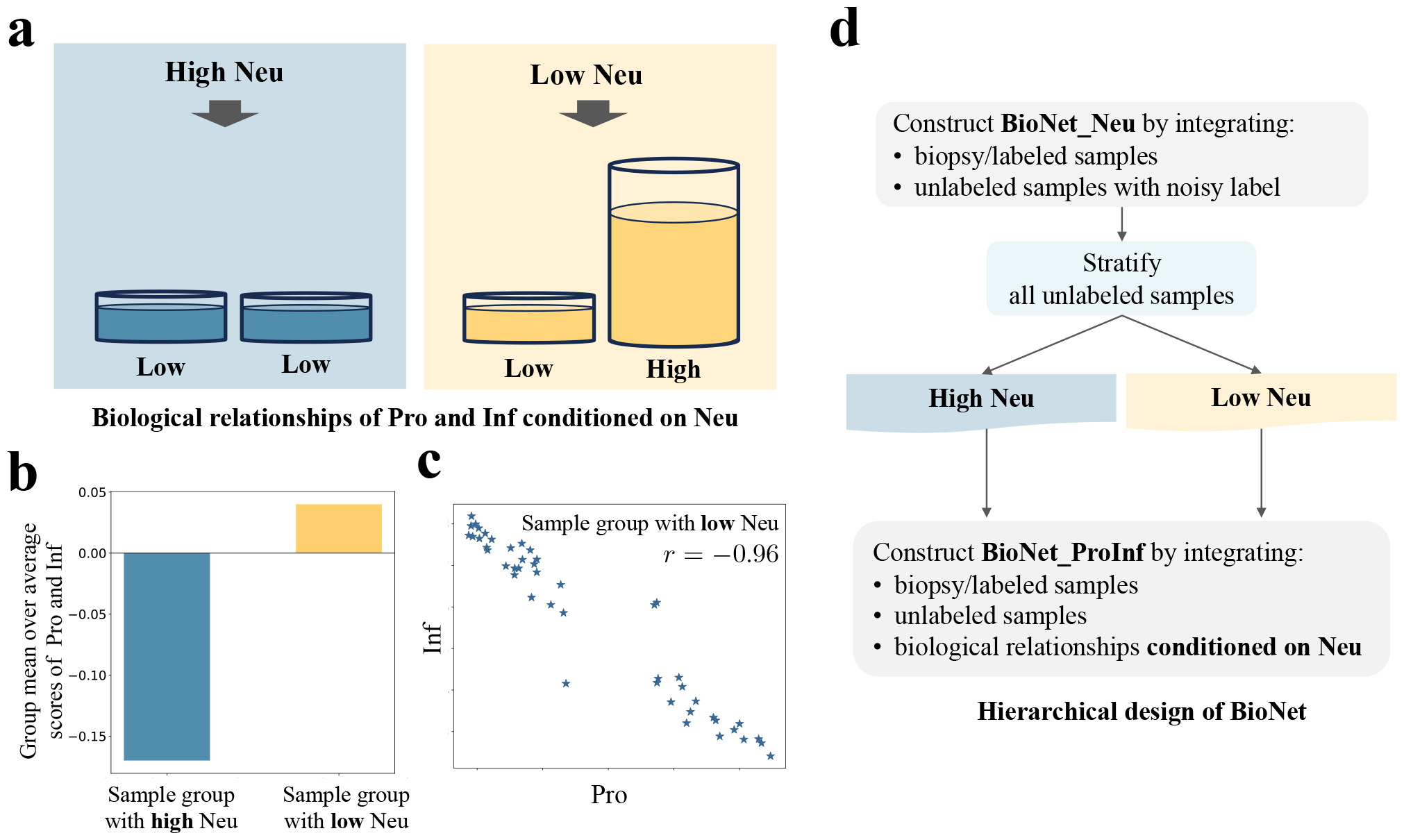
**(a)** Revealing biological relationships from domain knowledge to inspire BioNet design. **(b)** Bar chart for the group mean over the average scores of Pro and Inf in the group with high Neu in comparison to that in the group with low Neu. **(c)** Scatter plot of Pro and Inf for the group with low Neu. **(d)** Hierarchical design of BioNet inspired by two biological relationships (1) and (2)between Pro and Inf given the high/low status of Neu.

The aforementioned relationships are evident in biopsy samples. Specifically, we divided biopsy samples into two groups based on the enrichment scores of Neu: above-zero (high) and below-zero (low). **Fig. 2b** shows that samples with high Neu have significantly lower average scores of Pro and Inf, compared to samples with low Neu. This empirical evidence confirmed the previously mentioned relationship (1). **Fig. 2c** shows that samples with low Neu have a significant negative correlation between Pro and Inf. This confirmed relationship (2). It is worth emphasizing that while these relationships were demonstrated using biopsy samples, they were presumed to exist in unlabeled samples too, because these relationships were supported by fundamental knowledge about the spatial landscape of neuropathological alterations in GBM (30).

Incorporation of these knowledge-based relationships leads to a hierarchical design of BioNet (**Fig. 2d**). First, we constructed a neural network (NN), BioNet_Neu, to predict the Neu score using MRI. This model allowed us to stratify the large number of unlabeled samples into two groups with high and low predicted Neu scores, denoted as {*i* ∈ *Neu*^+^} and {*i* ∈ *Neu*^-^}, respectively. Next, we constructed another NN, BioNet_ProInf, a multitask learning architecture to simultaneously predict Pro and Inf using MRI. To deepen the understanding of the latent relationships between MRI features and the two primary gene modules, Pro and Inf, we integrated a trivial task, Neu, into BioNet_ProInf as a regularization strategy. This method was strategically implemented to encourage the model to discern the latent patterns in the data more effectively. A knowledge attention loss was designed and integrated into BioNet_ProInf to incorporate domain knowledge implicitly. This approach enabled the model to align closely with established expert understanding, thereby enhancing its predictive accuracy and generalizability.

### Construction of BioNet_Neu to predict Neu using transfer learning and uncertainty quantification

The overall architecture of BioNet_Neu is presented in **Fig. 3**. We adopted several strategies to tackle the challenge of a small biopsy sample size: (1) Employed transfer learning by pre-training the network using a large number of unlabeled samples who have noisy Neu labels informed by biological knowledge, then fine-tuning it using real biopsy samples.

**Fig. 3.**
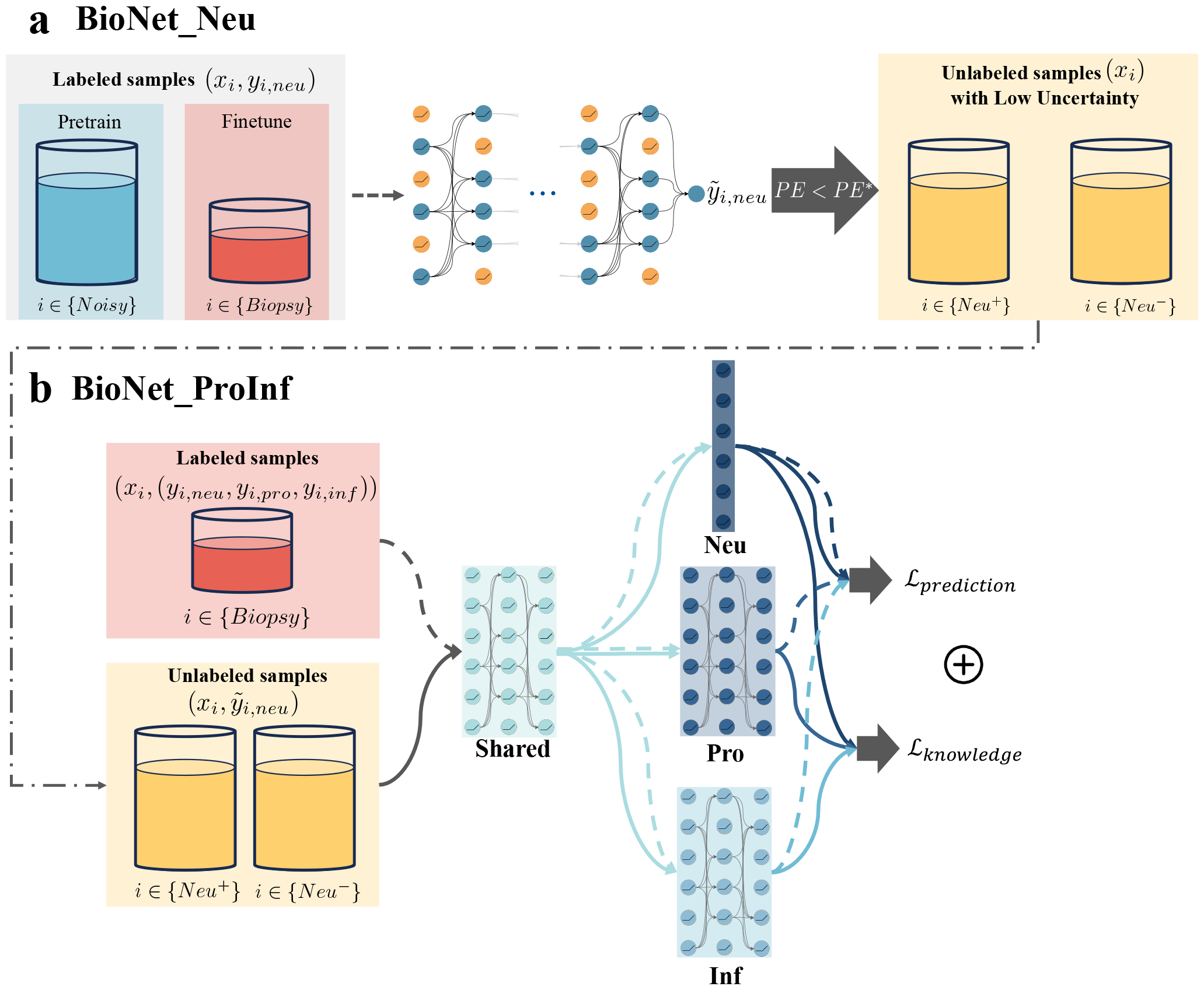
Overall architecture of BioNet. **BioNet** consists of two networks: BioNet_Neu to predict Neu using MRI; BioNet_ProInf to simultaneously predict Pro and Inf using MRI. **BioNet_Neu** is a feedforward neural network pre-trained using a large number of unlabeled samples with noisy Neu labels informed by biological knowledge, and fine-tuned using biopsy samples with data augmentation. It also corporates Monte Carlo dropout to enable uncertainty quantification for the predictions. The role of BioNet_Neu is to stratify unlabeled samples with high predictive certainty, which were then incorporated into the training of BioNet_ProInf. **BioNet_ProInf** is a multitask semi-supervised learning model with a custom loss function. The architecture consists of a shared block and task-specific blocks. The loss function combines a prediction loss and a knowledge attention loss that penalizes violation of the knowledge-based relationships on unlabeled samples.

(2) Incorporated Monte Carlo dropout (38) to enable uncertainty quantification (UQ) of the predictions. (3) Applied data augmentation by including neighboring samples of each biopsy sample in training.

In the hierarchical design of BioNet, BioNet_Neu played an important role in stratifying unlabeled samples into high and low predicted Neu groups. As the subsequent model was dependent on this sample stratification, we aimed to select unlabeled samples which had high predictive certainty. This highlighted the importance of the UQ capability of BioNet_Neu. To evaluate the UQ capability of a DL model, a common strategy is to examine if the model satisfies the “more certain more accurate (MCMA)” criterion (39), indicating that predictions with higher certainty are more accurate. To evaluate this criterion for BioNet_Neu, we first computed the predictive entropy (PE) (40) as an uncertainty score for each biopsy sample. We then computed the accuracy on subsets of samples above increasingly stringent PE thresholds. The accuracy increased from 71% to 90% when computed on the top certain samples. To ensure the selected samples have relatively high accuracy, we set a threshold, PE*, corresponding to a 90% accuracy level, and retained only those samples for which PE is less than PE*.

### Construction of BioNet_ProInf to predict Pro and Inf by combining multitask learning, semi-supervised learning, and domain knowledge

BioNet_ProInf is a multitask, semi-supervised learning model with a custom loss function. The input for BioNet_ProInf consists of biopsy/labeled samples, along with unlabeled samples that are selected and stratified by BioNet_Neu. The overall architecture of BioNet_ProInf is presented in **Fig. 3**. Domain knowledge indicates a pronounced relationship between Pro and Inf conditioned on their corresponding Neu. Consequently, the discriminative features identified for Neu are posited to be beneficial in the classification tasks for Pro and Inf. Inspired by this premise, we incorporated the auxiliary task Neu into the BioNet_ProInf architecture as a regularization strategy. Rather than explicitly providing the model with labels for Neu, we designed it to independently discern the relationships between MRI features and Neu. The linear classification layer of Neu (indicated in deep blue in **Fig. 3**) compels the model to embed discriminative features of Neu within the final shared layers’ outputs, thereby enriching the feature representation for two primary classification tasks. The loss function imposes penalties on prediction errors *ℒ*_*prediction*_ as well as violation of the knowledge-based relationships *ℒ*_*knowledge*_ Prediction errors for Pro and Inf are computed using biopsy/labeled samples. Conversely, prediction errors for Neu are derived from both labeled samples and unlabeled samples with high certainty predicted labels 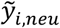 from BioNet_Neu. This approach strengthens the generalization of discriminative features for Neu. Violations of relationships are characterized by a knowledge attention loss, and defined on unlabeled samples. To subtly incorporate Neu labels into the model, we propose using predicted labels *ŷ*_*I,neu*_ from BioNet_ProInf for calculating the knowledge loss, rather than the true labels. This strategy stems from the observation that learning the relationships between Neu and the primary gene modules, Pro and Inf, is relatively straightforward. In contrast, mapping MRI features to these gene modules is significantly more complex and challenging. Employing true labels in the knowledge loss could lead the model to prioritize learning the simpler task and neglect the more critical yet difficult ones. Consequently, after perfectly learning the easier tasks, the model starts to predict Pro and Inf based solely on Neu, disregarding the MRI features. This is a scenario we aim to avoid, as it could undermine the model’s ability to truly understand and leverage the intricate relationships between MRI features and the gene modules. There are three components in *L*_*knowledge*_. The first two are tailored to correspond with two knowledge-based relationships, for Neu high and Neu low respectively. The third component is a barrier loss (41) defined on all unlabeled samples, aiming to discourage the predicted *ŷ*_*I,pro*_ and *ŷ*_*I,inf*_ from both being high.

We compared BioNet_ProInf with a range of existing models: (1) Supervised learning models such as feed-forward neural network (NN), support vector regression (SVR), and random forest (RF), which used only biopsy/labeled samples; (2) Semi-supervised learning (SSL) that utilized both labeled and unlabeled samples. We included a recent method called AdaMatch (42), a refined model for the highly-cited FixMatch (43). (3) Multitask learning (MTL) that exploited the relationship between multiple outputs such as Pro and Inf in our case. We included a supervised MTL-NN and a recent semi-supervised MTL model called multitask adversarial autoencoder (MTL-AAE) (44). The hyperparameters of BioNet and all competing methods were systematically tuned based on the same data-driven criterion through random search and grid search techniques. The details about hyperparameter tuning can be found in **SI Sec. 6** and **Sec. 7**.

As shown in **Fig. 4a**, BioNet_ProInf achieved AUCs of 0.80 and 0.81 for predicting Pro and Inf, respectively, on cohort A using leave-one-patient-out CV (LOPO CV). The classification accuracies (ACCs) by dichotomizing the scores into high and low were 80% and 75%. In contrast, despite extensive hyperparameter tuning, competing methods demonstrated limitations, achieving only modest performance metrics: AUCs lower or around 0.6 and ACCs lower or around 70%. This outcome not only underscores the intrinsic challenges posed by the task but also reflects the complexity of the underlying data. Presently, even experienced experts face difficulties in differentiating between Pro and Inf. Similarly, current data-driven approaches fall short in accurately classifying these two gene states, indicating a significant gap in the predictive capability of existing models.

**Fig. 4.**
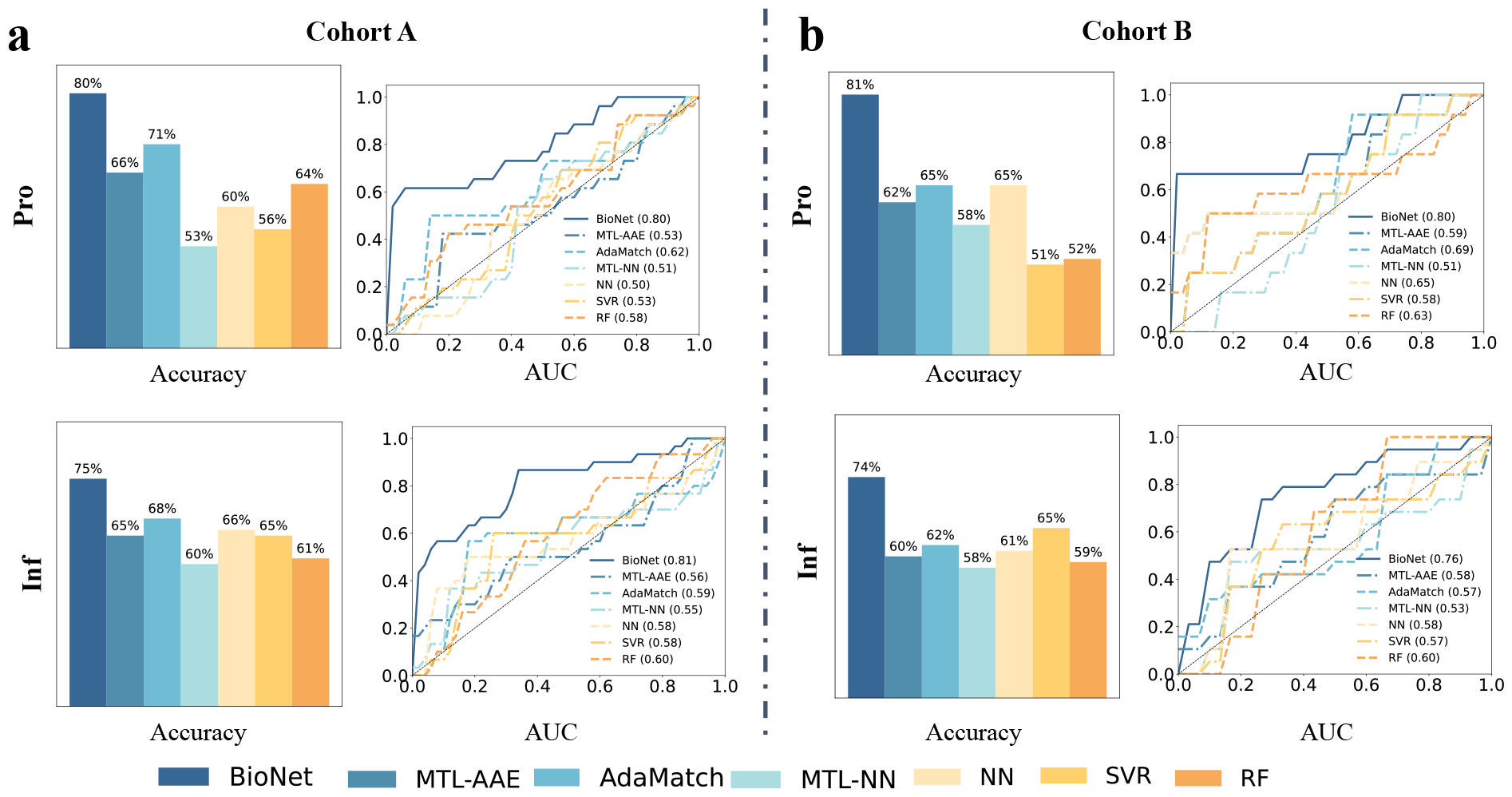
Performance of BioNet and competing methods on **(a)** developmental cohort A under leave-one-patient-out cross validation(LOPO CV) and **(b)** test cohort B.

### Testing of BioNet in an independent cohort B

The primary objective of our study was to evaluate the effectiveness of BioNet in its capability to precisely predict the Pro and Inf gene states in unseen datasets. This assessment aimed to establish the model’s generalizability across different datasets, underscoring its potential applicability in broader gene state classification tasks. As shown in **Fig. 4b**, BioNet achieved AUCs of 0.80 and 0.76 for predicting Pro and Inf, respectively, on cohort B. The ACCs by dichotomizing the scores into high and low were 81% and 74%. In comparison, the competing methods achieved AUCs ranging in 0.51-0.69 and 0.53-0.60, ACCs ranging in 51% -65% and 58% - 65%, for predicting Pro and Inf. BioNet outperformed all the competing methods.

### Evaluating the generalizability of BioNet on both cohorts

The most robust assessment of model performance is derived from the prediction accuracy of biopsy samples, as previously indicated. However, due to their sparse nature, biopsy samples are not suitable for evaluating the generalizability of each method. To overcome this, BioNet_ProInf and its competing methods were utilized to predict the Pro and Inf scores for unlabeled samples within each AOI. To assess the models’ generalizability, we proposed to leverage two known relationships between gene modules. This analysis served as an effective method to validate the predictive accuracy of each model beyond the biopsy locations. Specifically, we computed two knowledge concordance (KC) metrics, denoted as *KC*_*neu*_^+^ and *KC*_*neu*_^-^, to quantify the concordance between the model’s predictions and the relationships. The results were presented in **Fig. 5b**.

**Fig. 5.**
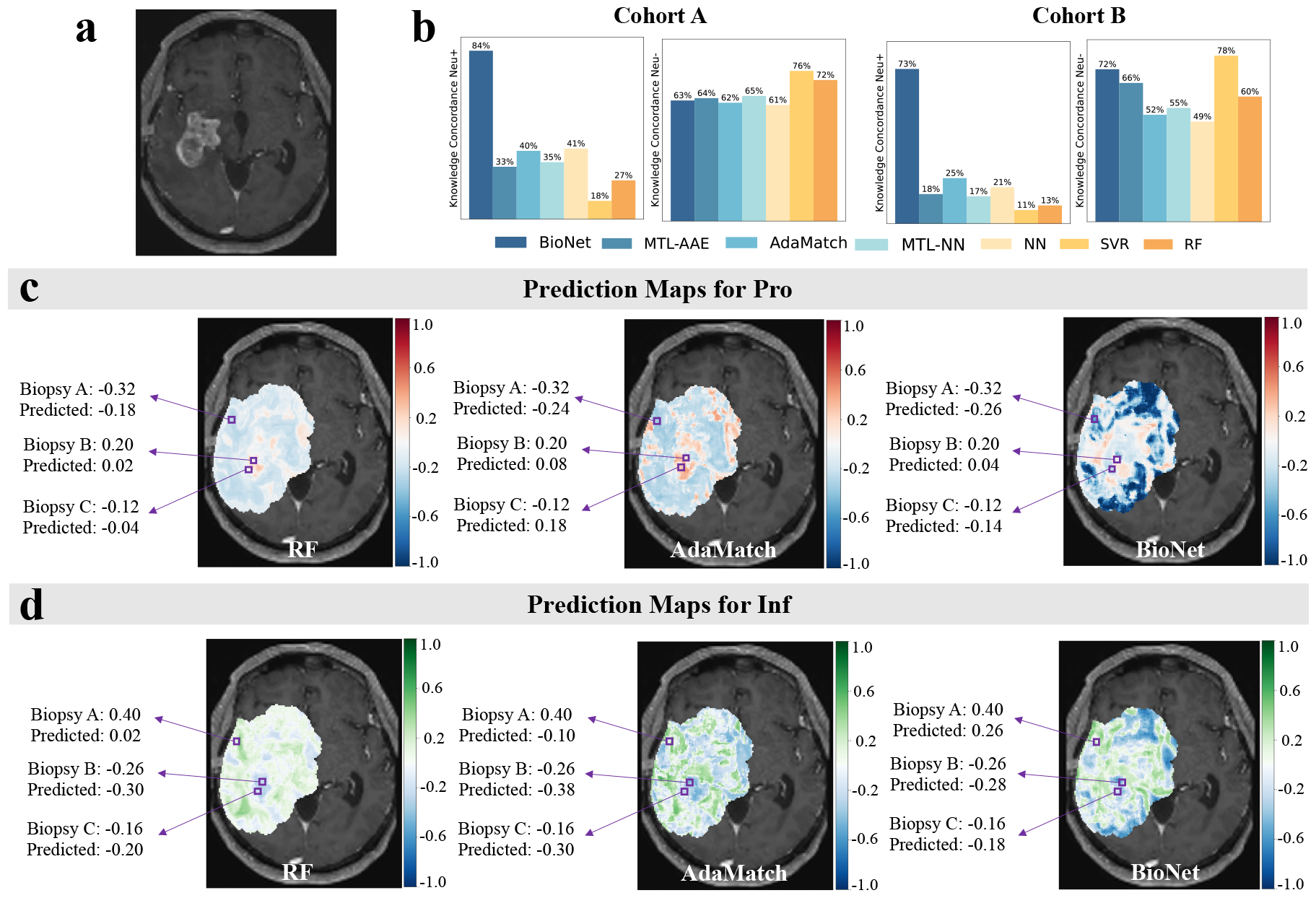
Generalizability of BioNet_ProInf and competing methods. **(a)** The patient’s T1Gd MRI image. **(b)** Knowledge concordance(KC) metrics for the predictions on unlabeled samples from the tumoral area of interest (AOI) of each patient on developmental cohort A and test cohort B. Prediction maps of **(c)** recGBM_Pro and **(d)** recGBM_Inf within the tumoral AOI by BioNet and the best-performing ML and DL methods (color maps overlaid on **(a)**). Three purple boxes denote the locations of three biopsies on this MRI slice. The maps indicate that BioNet attains the lowest absolute errors in predicting both Pro and Inf.

BioNet_ProInf demonstrated moderately high KC metrics, exceeding 60% in both cohorts. In contrast, the *KC*_*neu*_^+^ values for all competing methods did not surpass 50% in either cohort. These values suggest that the incorporation of knowledge regularization into these models was effective. However, it is important to recognize the inherent uncertainty in domain knowledge. Consequently, overly strong regularization, potentially leading to very high concordance, was not desirable. Instead, we aimed for a balanced approach where models harmonize knowledge regularization with prediction losses. From this perspective, achieving moderately high concordance aligns well with our objectives. Furthermore, the observation that all methods demonstrated moderately high *KC*_*neu*_^-^suggests that this relationship might be inherently more learnable for models, even without explicit regularization.

To provide a visualization for the intratumoral heterogeneity of gene modules across the AOI, prediction maps of two gene modules were generated for each patient. **Fig. 5** displays the prediction maps on a selected MRI slice, specifically chosen for having the most biopsies in a single slice. The prediction maps in **Fig. 5cd** were from BioNet and the best-performing ML and DL competing methods (color maps overlaid on the patient’s T1Gd MRI image **Fig. 5a**). The majority of locations in the prediction maps generated by competing methods, especially by RF, are with light color, which suggests a tendency of these models to predict most samples with gene module scores around zero. Such a pattern indicates a degree of uncertainty in the classification, reflecting a potential limitation in the models’ discriminative capabilities. Supporting this, the average prediction entropy for BioNet is 0.588, markedly lower than that of RF (0.683) and AdaMatch (0.667). Compared with competing methods, BioNet stands out by delivering not only more accurate predictions on biopsies but also exhibiting significantly higher confidence in its predictions on unlabeled samples.

## Discussion

In this study, we utilized a unique dataset and developed BioNet to facilitate non-invasive quantification of intratumoral heterogeneity of gene modules in recGBM patients. The differentiation of proliferative/recurrent tumor (Pro) and treatment-induced reactive/inflammatory cells (Inf) is crucial for evaluating treatment effectiveness and guiding subsequent therapeutic strategies, but is very challenging in clinical practice due to their indistinguishable appearances on MRI. Biopsy, as the gold-standard approach, is invasive and only samples a few sparse regions. Our approach enabled the mapping of the regional distribution of these gene modules across the entire lesion for each patient. It enhances the accuracy of surgical targeting for confirmatory biopsies, assists in evaluating treatment effectiveness, enables regular monitoring of the gene modules for timely identification of treatment response or resistance, and provides a deeper understanding of patient-specific nuances to tailor treatment more effectively.

This study has several limitations. One limitation of this study is the constrained sample size of biopsy samples in both cohorts. Although BioNet is innovatively designed to mitigate this limitation by leveraging unlabeled samples and domain knowledge, the pivotal role of biopsy/labeled samples in training an accurate and robust predictive model cannot be overstated. Increasing the number of biopsy samples has the potential to significantly improve the prediction accuracy of BioNet. Nevertheless, the invasive nature of biopsies restricts the feasibility of acquiring large sample sizes from individual patients. Future research should therefore aim to expand patient enrollment across multiple centers, facilitating a more substantial collective sample base. Additionally, considering recent findings on sex differences in GBM (45), the development of demographic-specific models, such as sex-specific variants, may further refine prediction accuracy. The second bottleneck of this work is the model’s explainability. Although BioNet has demonstrated effectiveness in predicting regional distributions of gene modules using MRI, the complexity of the model poses challenges in understanding the underlying mechanisms driving the predictions. The complex architecture and numerous parameters of DL models frequently result in a lack of interpretability. Future research should therefore prioritize enhancing the interpretability and explainability of BioNet. Investigating methodologies such as advanced visualization tools, and model-agnostic interpretability approaches could provide valuable insights into the specific image features and patterns that BioNet leverages for its predictions. Such advancements will not only guarantee that decisions are made based on accurate and comprehensible information, but will also offer critical insights into the growth of different gene modules, thereby paving the way for future research in GBM.

## Methods

### Patient inclusion and biopsy acquisition

This study involved a developmental cohort (A) and a test cohort (B). Cohort A was acquired as part of a retrospective, observational study designed to study patients who had undergone repeat surgical resection for recurrence of high-grade gliomas following chemotherapy and radiation therapy (30). Cohort B was acquired as part of a prospective, clinical trial for convection enhanced delivery of topotecan chemotherapy (46). Both cohorts received standard of care neuroimaging performed at Columbia University Irving Medical Center within one month prior to surgery (Cohort A) or one day prior to surgery (Cohort B). The neuroimaging exam of each patient produced multiparametric MRI data such as T1-weighted+Gd (T1Gd), T2-weighted (T2), FLAIR, apparent diffusion coefficient (ADC), and susceptibility weighted imaging (SWI). Biopsy samples from each patient were obtained according to IRB-approved protocol from the operating room (see details in **Supplementary Information (SI) Sec. 1**). Patient information was de-identified and maintained by a tissue broker who has designated clinical information. Specifically, cohort A included 84 biopsies harvested from 37 patients (mean age = 56, 63% male, 1-3 biopsies per patient); cohort B included 31 biopsies from 5 patients (median age = 56, 60% male, 1-10 biopsies per patient).

### RNAseq-IHC correlation analysis, clustering, and GSVA

For samples in Cohort A that had both RNAseq and IHC quantification, Pearson correlation was calculated between the normalized expression values for each gene and the IHC labeling index for each marker, building a correlation matrix of 15001 genes by 5 IHC markers (4 IHC markers and total normalized cellularity from H&E images). The 15001 genes were filtered from 23802 genes to include only protein coding genes that had more than 10 reads across all samples. A p-value cutoff of 0.05 (un-adjusted) was set for determining the significance of the IHC-gene expression correlation for each stain, and all genes with significant correlations with ≥1 marker were selected for downstream analysis. Furthermore, based on the resulting correlation matrix, we performed clustering analysis. A variety of different clustering algorithms were used, which repeatedly found the optimal cluster number to be two or three. Considering a recent publication by our group that described a three-tissue-state model of glioma (30), we ultimately decided to proceed with the three-cluster result from hierarchical clustering with Ward’s linkage. Three major clusters with mutually exclusive genes were identified, and these gene clusters were used as gene sets for downstream Gene Set Variation Analysis (GSVA) on a sample-by-sample basis, which produced three gene set enrichment scores for each MRI-localized biopsy. We call the three gene sets “gene modules” and use the enrichment scores as estimates of gene module expression. Given that there exists a degree of noise in bulk RNAseq analysis, this technique allows us to leverage the multi-modal data (RNAseq and IHC) to link expression with biologically meaningful tissue features (IHC markers) to further strengthen the gene module approach. Analysis of the genes in each module suggested that the modules consisted of proliferative, inflammatory, and neuronal genes. Thus, the modules were named Pro, Neu, and Inf in this paper. The enrichment scores of these modules ranged from -1 to 1, and were linearly transformed to [0,1] during BioNet training. For a detailed description of the RNA extraction and pooled library amplification for transcriptome expression RNA-sequencing (PLATE-Seq), please refer to **SI Sec. 2**. For a detailed description of IHC and quantification, please refer to **SI Sec. 3**.

### Identification of three tissue-specific gene modules: Neu, Pro, Inf

To reduce complexity and improve prediction accuracy, we amplified the signal-to-noise ratio of genetic and cellular heterogeneity signal in the tissue by combining individual gene expressions and cellular composition patterns into three *gene modules*. While the radiogenomic signal associated with individual genes can be noisy, there is a great potential to improve the accuracy of ML/DL models by targeting on predicting clusters of correlated genes that represent different gene modules. This analysis was performed on cohort A. The majority of biopsies (48/84; 57%) had both RNA sequencing and IHC staining for SOX2, CD68, Ki67, and NeuN, where SOX2 is described as a robust marker for glioma cells (47); CD68 is a known marker of macrophages; Ki67 is a known marker of proliferation; NeuN, also known as RBFOX3, is a canonical marker of neurons. These markers were used as (non-comprehensive) proxies for different cell populations in the glioma microenvironment. Hemotoxylin counterstain was used to label all nuclei, providing a measure of total cell density. As a first step to “pre-screen” genes for our downstream analysis, we performed correlation analysis between normalized gene expression values and IHC stained cell counts, and identified 7779 statistically significant genes (p-value < 0.05, un-adjusted). Hierarchical clustering determined three distinct clusters/modules with mutually exclusive genes **(Fig. 6a)**. Module 1-3 consisted of 3688, 1673, and 2418 genes, which have significant positive correlations with the labeling indices of SOX2, Ki67, and total cell density, the labeling index of NeuN, and the labeling index of CD68, respectively. For a complete list of the genes included in each module, please refer to **Supplemental Table S1** in SI. Gene ontology analysis demonstrated that module 1-3 were associated with proliferative (Pro) -proliferation/cell cycle ontologies; inflammatory (Inf) -cytokine production/immune response; neuronal (Neu) - neuronal signaling, respectively (**Fig. 6b**). These gene modules developed from the RNA-IHC clustering were then applied to all 84 biopsies via Gene Set Variation Analysis (GSVA), using the normalized PLATE-seq expression profile for each sample (**Fig. 6c**). The genes identified in cohort A were used to perform GSVA on cohort B. In this paper, the gene modules were named Pro, Inf, and Neu, whose values are the GSVA enrichment scores. In this study, the raw enrichment scores of three gene modules, originally ranging from -1 to 1, are transformed to a [0, 1] scale using the mapping function 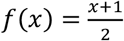 Consequently, the threshold distinguishing high and low gene module expression is correspondingly transformed from 0 to 0.5.

**Fig. 6.**
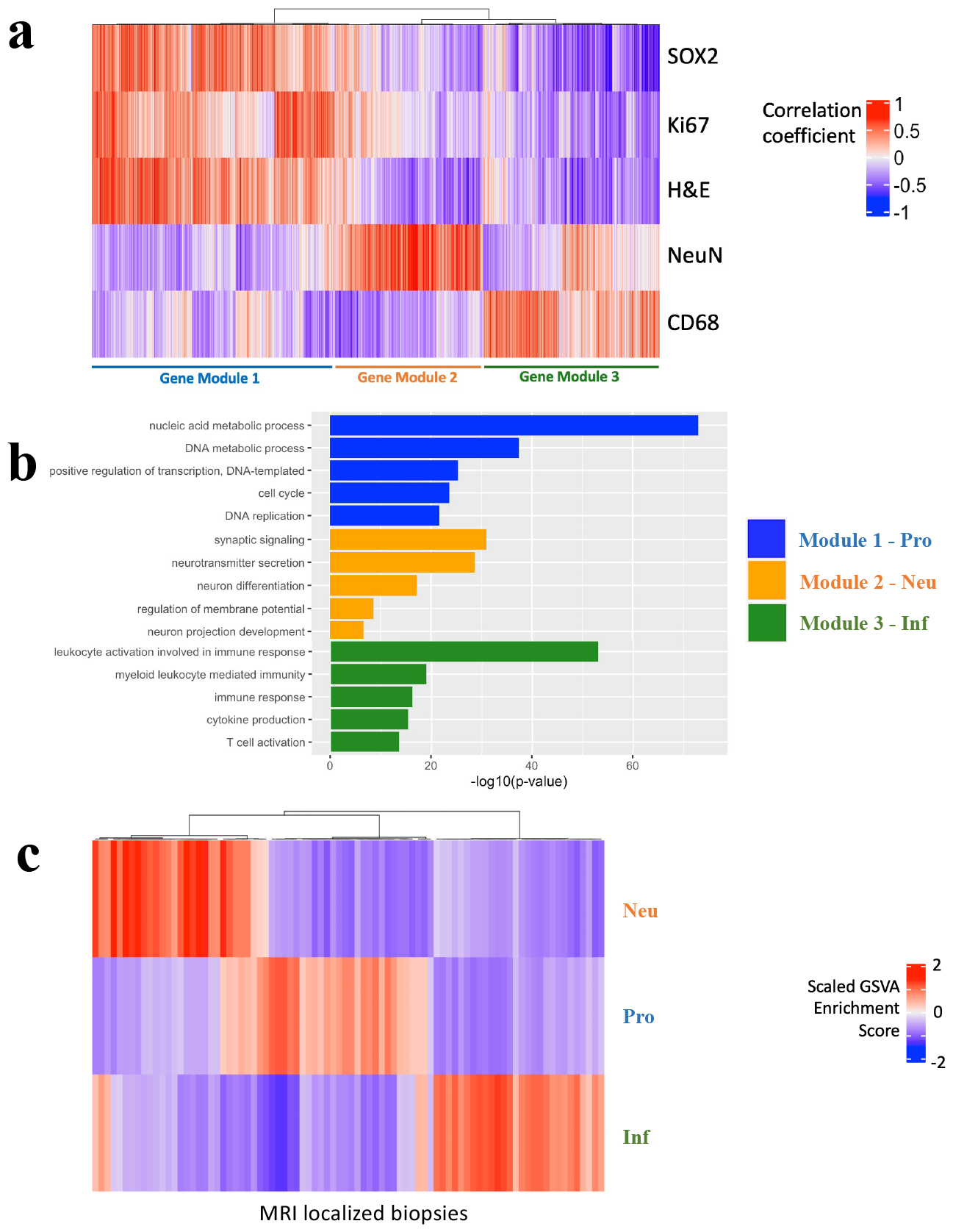
Defining tissue-specific gene modules to connect with key immunohistochemical features. **(a)** Heatmap depicting correlation between normalized gene expression and immunohistochemical labeling indices, with subsequent hierarchical clustering revealed three orthogonal tissue-specific gene modules. Module 1 consists of 3688 genes significantly positively correlated with SOX2/Ki67/H&E; module 2 consists of 1673 genes correlated with NeuN; module 3 consists of 2418 genes correlated with CD68. **(b)** Bar plot depicting top significant gene ontologies enriched in each of the three tissue-specific gene modules derived from the IHC-RNAseq correlation analysis. X-axis is –log_10_(p-value) of each ontology. A vertical dash line was placed at p-value = 0.05. Module 1 is enriched in genes involved in proliferation (Pro), module 2 in neuronal-specific genes (Neu), and module 3 in genes in immune infiltration (Inf). **(c)** Heatmap depicting single-sample GSVA analysis for each of 84 MRI-localized biopsies for each of the three tissue-specific gene modules. Color gradient represents magnitude/direction of tissue-specific enrichment for each biopsy.

### Extraction of regional features from multiparametric MRI

The regions were defined as 5×5 pixel^2^ windows on the axial view of MRI. This size approximated the physical size of biopsy samples. For each biopsy sample, image features were extracted from a window at the biopsy location. This resulted in a labeled dataset 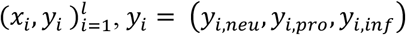, with 69 samples corresponding to biopsies obtained from 31 patients in cohort A (1-3 per patient). This was a subset of the total 84 samples due to file corruption or missing scans. Additionally, image features were extracted from regions beyond the biopsy locations within a tumoral AOI, to generate unlabeled samples. To do this, we first defined the AOI of each patient by combining the segmented enhancing tumor portion on T1Gd and infiltrating tumor portion on T2/FLAIR plus a 7mm margin. Then, we placed sliding windows with a size of 5×5 pixel^2^ and a stride size of 1 throughout the AOI, and extract image features from each window (**Fig. 1** Texture feature extraction). This resulted in an unlabeled dataset 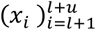 with about 1.82e6 samples (5e3 to 9e4 per patient). The regional image feature set consisted of 280 statistical and texture features computed from T1Gd, T2, FLAIR, ADC and SWI. For a detailed description of MRI preprocessing, segmentation, and feature extraction, please refer to **SI Sec. 4**.

### Construction of BioNet using cohort A

BioNet includes two networks: 1) BioNet_Neu is to predict Neu; 2) BioNet_ProInf is to simultaneously predict Pro and Inf.

### 1. BioNet_Neu

BioNet_Neu achieved an Area Under the Curve (AUC) of 0.77 based on Cohort A using 5-fold cross validation (CV). Without Monte Carlo dropout, transfer learning and data augmentation, the AUC was reduced to 0.70, 0.64 and 0.56. In the data augmentation approach, including samples within a 5-voxel radius of the biopsy sample yielded the most optimal performance. Additional details can be found in **SI Sec. 5**.

#### Architecture

This network included two 2048-dimension hidden layers with ReLU as the activation function. To incorporate UQ, Monte Carlo dropout was adopted with a dropout rate of 1e-3.

#### Pre-training

We created a large, noisy labeled dataset, denoted as {*Noisy*}, to pre-train the network. The noisy labeled dataset consisted of unlabeled samples that were likely to have high or low Neu based on domain knowledge, denoted as class 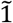 or class 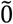, respectively. The overhead ‘∼’ indicates that the knowledge has uncertainty, which may result in labeling errors of the samples. However, since this dataset was solely used for pre-training purposes, such errors were deemed acceptable.

In detail, we have the knowledge that Neu tends to be high on the boundary of AOI and outside the AOI (i.e., in the normal brain areas), as the genes included Neu are predominantly involved in neuronal signaling. To avoid unwanted brain structures, we chose to include samples located on the AOI boundary as class 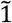 samples. Furthermore, we have the knowledge that Neu tends to be low within the enhancing tumoral area on T1Gd, as this region is known to involve tumor proliferation or immune response (48) rather than neuronal signaling. Thus, we included samples from the enhancing tumoral area as class 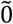 samples. As a result, the noisy labeled dataset included about 7500 samples in class 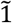 and 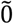, respectively, from 31 patients. Using this dataset, we pre-trained the network under the cross-entropy loss to differentiate between class 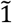 and 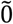.

#### Fine-tuning

The pre-trained network was fined-tuned using biopsy samples under the soft cross-entropy loss, considering that scaled Neu scores ranged between 0 and 1. Data augmentation was used by incorporating neighbor samples within a 5-pixel radius of each biopsy sample.

#### Prediction and stratification of unlabeled samples

The network was used to predict the Neu scores for unlabeled samples within the ROI of each patient. Recall that the unlabeled samples corresponded to 5×5 pixel^2^ windows with a stride of 1, sliding over the AOI. Using zero as a cutoff, the predicted scores were dichotomized into two classes,{*Neu*^+^} or {*Neu*^-^}. Furthermore, the network’s UQ capability made it possible to generate an uncertainty score for each prediction, measured by Predictive Entropy (PE) (40). To filter out unlabeled samples whose predictions have high certainty, we applied a threshold PE**, corresponding to a 90% accuracy level, and only retained samples with PE<PE*. The retained samples were then divided into two subsets: {*i* ∈ *Neu*^+^} and {*i* ∈ *Neu*^-^}, which included unlabeled samples predicted to be *Neu*^+^ or *Neu*^-)^ with high certainty, respectively. These subsets would be used in training BioNet_ProInf as discussed in the following section.

### 2. BioNet_ProInf

#### Architecture

the network used to predict Pro and Inf is a multitask semi-supervised learning model with a custom loss function. It comprises a shared block and task-specific blocks. The shared block consists of three layers with dimensions of 256, 128, and 128, respectively. The task-specific blocks for Pro and Inf each consist of three layers with dimensions of 128, 128, and 64, respectively. As the relationships between Pro and Inf are conditional on the status of Neu, discriminant features of Neu provided significant guidance for the main tasks. To enforce the shared latent representations to encode the discriminant features, the auxiliary task block corresponding to Neu was kept simple, comprising one layer of 128 units. Input to the network included both biopsy/labeled samples and unlabeled samples in {*i* ∈ *Neu*^+^}U{*i* ∈ *Neu*^-^} which were selected by BioNet_Neu as previously described. The output was used to define a custom loss function, introduced as follows:

#### Loss function

There are two parts in the loss function to penalize (1) prediction errors on biopsy/labeled samples, and (2) violation of domain knowledge, i.e.,

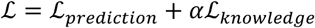

To compute *ℒ*_*prediction*_, the network generated three predicted scores of Pro, Inf, Neu for each biopsy sample, which were compared with the true scores using the L2 norm for two main tasks and Kullback−Leibler divergence for the auxiliary task. In addition, prediction errors for Neu are also derived from unlabeled samples with high certainty predicted labels 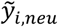 from BioNet_Neu, i.e.,

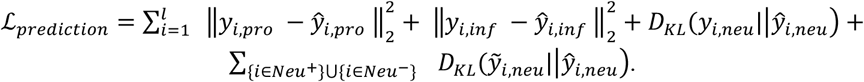

The knowledge attention loss *ℒ*_*knowledge*_ is defined on unlabeled samples. To implicitly incorporate the labels for Neu into the model, we utilize the predicted labels *ŷ*_*I,neu*_ in *ℒ*_*knowledge*_, which is designed to include three components:

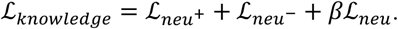

Here,

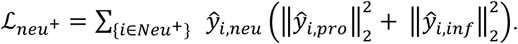

Minimizing this loss encourages the predicted *ŷ*_*I,pro*_ and *ŷ*_*I,inf*_ to be low for unlabeled samples in {*i* ∈ *Neu*^+^}, where *ŷ*_*I,neu*_ acts as a weight, promoting a higher occurrence of such predictions for samples with higher *ŷ*_*I,neu*._

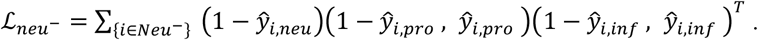

Minimizing this loss encourages the predicted *ŷ*_*I,pro*_ and *ŷ*_*I,inf*_ to be negatively correlated for unlabeled samples in{*i* ∈ *Neu*^-^}, where (1*− ŷ*_*I,neu*_) acts as a weight, promoting a higher occurrence of such predictions for samples with lower *ŷ*_*I,neu*_.

*ℒ*_neu_ is a barrier loss (41) defined on all unlabeled samples {*i* ∈ *Neu*^+^}U{*i* ∈ *Neu*^-^}, aiming to discourage the predicted *ŷ*_*i,pro*_ and *ŷ*_*I,inf*_ from both being high. This loss can help strengthen the effect of the other two losses, with its utility controlled by *β*. The form of *ℒ*_neu_ follows the standard log barrier function commonly found in optimization literature, i.e.,

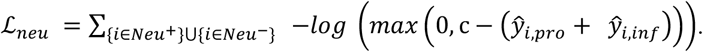

c is an upper bound that can be treated as a tuning parameter. However, for simplicity of the design, we set c=1.2, which is the maximum value of the summation of scaled Pro and Inf scores in biopsy samples.

### Testing of BioNet using cohort B

The MRI scans in cohort B were acquired at a different resolution compared to cohort A. Using this test cohort could provide valuable insights into the generalizability of BioNet on a less ideal yet more realistic dataset, reflecting the common practice that MRI scans can be obtained under varying conditions for different patients. However, the resolution discrepancy created challenges in directly applying the trained BioNet from cohort A to cohort B. In DL, approaches that address input discrepancy when deploying a model from one domain to another domain have been explored in the subfield of domain adaptation (49).

To address this discrepancy, we implemented an approach inspired by domain adaptation, which replaced the unlabeled samples from cohort A with those from cohort B to re-train BioNet_ProInf. The unlabeled samples were abundant and contained only image features. Using the unlabeled samples from cohort B had the effect of biasing BioNet_ProInf toward the image representation in cohort B. Notably, this re-training process did not include any biopsy sample from cohort B. Thus, the re-trained BioNet_ProInf was still “blind” to the ground-truth scores of the biopsy samples in cohort B.

## Acknowledgement

This work was supported by NIH grant U01CA250481-01A1, NSF grant DMS-2053170, NIH/NINDS R01NS103473, NIH/NCI R01CA161404 and the NIH/NCI Cancer Center Support Grant P30CA013696. We would like to acknowledge the Herbert Irving Comprehensive Cancer Center Molecular and Pathology Core.

## Data availability

The datasets for Cohort A and Cohort B are accessible on Figshare through the project page: https://figshare.com/projects/Texture_features_of_Multiparametric_MRI_-_Recurrent_Glioblastoma/193223, with the dataset available at the following DOI: https://doi.org/10.6084/m9.figshare.23950584.v1. RNAseq and enrichment analysis of these datasets have been published in (30).

## Code availability

The code used to extract texture features, perform experiments, and analyse data is available at: https://github.com/hairongw/BioNet.git.

