## Supplementary Information for "Biologically-informed deep neural networks provide quantitative assessment of intratumoral heterogeneity in post-treatment glioblastoma"

### Supporting Information Text

#### 1. Biopsy acquisition

Fresh samples were divided into two pieces by trained neuropathologists and technicians. One piece was embedded in paraffin and underwent traditional H&E staining, while the second piece was stored for later use in a -80°C freezer. Intraoperatively, the neurosurgeon used the Brainlab neuro-navigation tool to identify the radiographic location of each biopsy. Biopsies were obtained from both contrast enhancing and contrast-negative, FLAIR-positive regions of tumor. A screenshot of the Brainlab localization was taken and custom registration software written in MATLAB (1) was later used to acquire the coordinates of the specific biopsy on the T1-post-contrast MRI image.

#### 2. RNA extraction and pooled library amplification for transcriptome expression RNA-sequencing (PLATE-Seq)

Total RNA was extracted from these tissue samples using the institutional genomic core. Samples with an RNA Integrity Number (RIN) greater than six were selected and subsequently sequenced using the PLATE-Seq protocol. PLATE-Seq is a novel sequencing platform that can produce high-throughput RNA-sequencing data through barcoding and pooling of cDNA libraries10. RNA samples from 84 MRI-localized samples were normalized to between 60-100 nanograms in 16.5 microliters of nuclease free water and placed in a 96-well PCR plate. Purified mRNA was reverse transcribed and barcode segments were added to create easily identified cDNA. Multiple batches were pooled, cDNA was purified, and then underwent PCR amplification. Samples were ultimately sequenced on an Illumina NextSeq 500 sequencer10. Raw reads were mapped to the human transcriptome by the institutional sequencing center using STAR alignment to generate gene counts. All RNAseq processing and analysis was performed using the statistical computing software R. RNA sequencing counts were first trimmed to only include genes with ≥10 counts across all samples. Counts were then normalized using the DESeq2 package to generate normalized expression values for each gene.

#### 3. Immunohistochemical staining (IHC) and quantification

A subset of the biopsies in Cohort A (n=48) and all biopsies in Cohort B underwent histological staining for H&E and IHC for SOX2, Ki67, CD68, and NeuN. Five-micrometer sections from a single localized biopsy were obtained for staining with hematoxylin-eosin and immunostaining with SOX2, Ki67, CD68, and NeuN. Slides were subsequently scanned and digitized at 40x magnification using a Leica SCN400 system (Leica Biosystems, Buffalo Grove, Illinois). Total cell density was calculated using a validated semi-automated whole slide cell-counting algorithm (2).

In brief, the algorithm was trained to select all hematoxylin-stained nuclei using 9 randomly generated high-power fields (HPF) from each sample, defined as .225 mm × .225 mm (900 × 900 pixels). Subsequently, the algorithm could be used to iteratively process all HPFs until the entire H&E stained slide was counted. After collecting the total cell counts for an H&E-stained slide, the algorithm was trained to select SOX2-stained and Ki67-stained nuclei on 9 randomly generated HPFs from SOX2-stained or Ki67-stained slides, respectively. For CD68-stained and NeuN-stained samples, the algorithm was trained to select CD68-stained and NeuN-stained cells in 9 randomly generated HPFs from CD68-stained or NeuN-stained slides, respectively. Areas of the slide without tissue were excluded from calculations in all cases. Algorithm-derived cell counts were manually verified and total cell density, SOX2 cell density, Ku67 cell density, CD68 cell density, and NeuN cell density were calculated across all HPFs in a slide. The mean cell count per HPF was used as a representative measure in all cases given that intra-specimen heterogeneity was limited by the standardized size of each biopsy. A labeling index (LI) for each immunostain was computed by dividing the number of immunostain-positive cells by the total cell count in a HPF (from the same slide). For H&E-stained slides, cell counts were converted to a percentage by dividing each by the largest cell count across all samples. Algorithm-derived cell counts were manually verified by a human reviewer. For validation, 100 HPFs were chosen at random and each field as manually inspected to determine the number of nuclei present. The same field was then evaluated by the automated cell-counting algorithm. A high correlation was observed between the automated algorithm and the manual cell counts for SOX2 and Ki67 cell density as determined by a Pearson coefficient, as shown in **Fig. S1**.

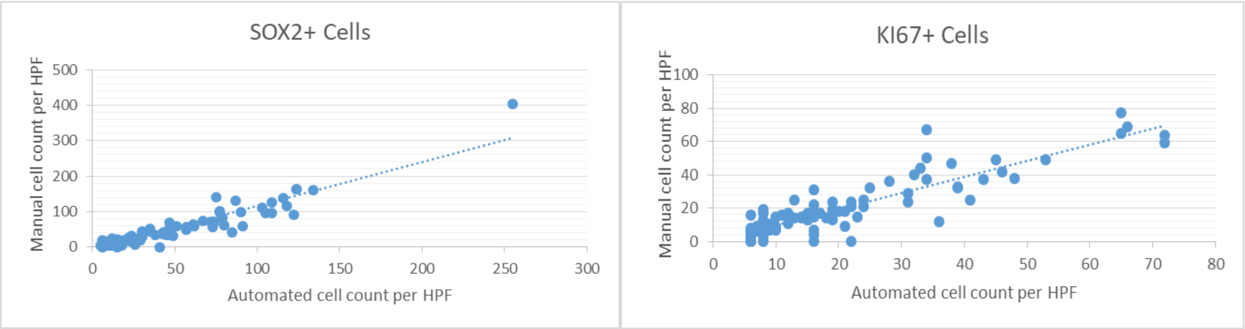


**Fig. S1.** Sample correlation between manual and automated cell counts (per HPF) for SOX2- and Ki67-stained nuclei, (r = .8917, r = .8123, respectively).

### 4. Multiparametric MRI pre-processing, segmentation, and feature extraction

Each patient received an MRI exam, which included T1Gd, T2, FLAIR, ADC, and SWI sequences. Detailed MRI parameters can be found in **Supplemental Table S2**. All images were preprocessed using a pipeline built in Python. For each patient, the multiparametric image set was affine registered (12 degrees of freedom) to the ADC image using SimpleElastix (3). Registration transforms were then applied to the associated tumor segmentation masks for each image. Next, inhomogeneity correction was performed using the N4 algorithm (4) implemented in SimpleITK (5). In addition, a brain mask was extracted for each patient using MONSTR, a multi-contrast brain-stripping tool (6). The brain mask was used to normalize the image intensities to a mean of zero and standard deviation of one for all images except the computed ADC. Finally, all images and segmentations were resampled to a common voxel size of 1.05 mm x 1.05 mm for data consistency across patients.

For each patient, the enhancing tumor and infiltrating tumor were manually segmented based on T1Gd and T2/FLAIR images by trained individuals using our in-house thresholding-based software. A grouped segmentation was then determined by combining the T1Gd and T2/FLAIR segmentations. This group segmentation was then dilated by a margin of 7 mm to create our AOI.

A sliding window of 5×5 pixel$^2$ was placed at each pixel within the AOI. From each sliding window, we extracted textural features from the ADC, FLAIR, SWI, T1Gd, and T2. Specifically, two commonly used texture analysis algorithms, Gray-Level Co-occurrence Matrix (GLCM) and Gabor Filters, were used to generate a total of 38 textural features for each of the five MRI images, for a total of 190 features. In addition, 18 commonly used 1$^{st}$-order statistical features such as mean, standard deviation, and energy were extracted, for a total of 90 1$^{st}$-order statistical features from the five MRI images. Collectively, our texture analysis pipeline generated 280 features for each sliding window. These features were also used in our prior radiomic study of GBM and shown to be effective for capturing imaging phenotypic information correlative with genetic and histopathological characteristics of the tumor (7–9).

### 5. Ablation study for BioNet_Neu

BioNet_Neu achieved an Area Under the Curve (AUC) of 0.77 based on Cohort A using 5-fold cross validation (CV) (**Fig. S2a**). Without Monte Carlo dropout, transfer learning and data augmentation, the AUC was reduced to 0.70, 0.64 and 0.56, respectively. **Fig. S2b** shows that BioNet_Neu satisfied the MCMA criterion.

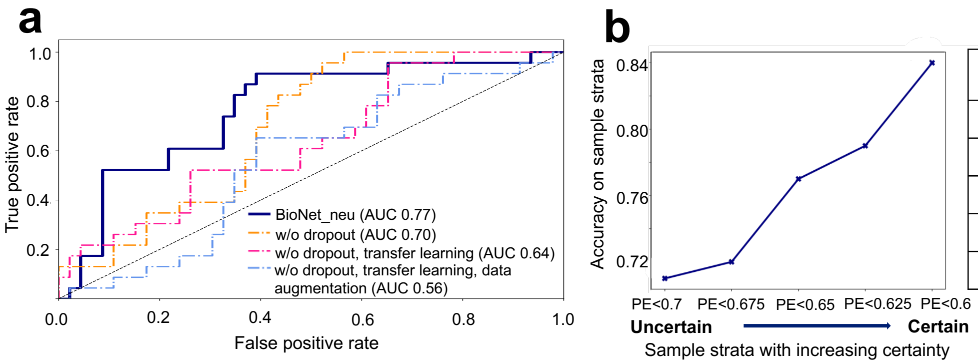


| Predictive entropy (PE) | Remaining samples | Accuracy |
|---|---|---|
| All | 100% | 71% |
| PE < 0.675 | 87% | 72% |
| PE < 0.650 | 72% | 77% |
| PE < 0.625 | 70% | 79% |
| PE < 0.600 | 59% | 84% |

**Fig. S2.** BioNet_Neu performance on cohort A. (**a**) ROCs of BioNet_Neu and ablated versions of the model. (**b**) Accuracy of BioNet_Neu evaluated on sample strata/subsets with increasing certainty.

#### 6. Hyperparameter tuning for BioNet_ProInf

There are two types of hyperparameters in BioNet_ProInf: generic and specific hyperparameters. Generic hyperparameters are common to most DL algorithms, including the number of layers, number of neurons per layer, learning rate, batch size, etc. Specific hyperparameters are unique to BioNet, being the weights used to combine multiple losses in the custom loss function, namely, $\alpha$ and $\beta$.

A grid search on all the hyperparameters is too time-consuming and prone to overfitting. Thus, we used a more efficient two-step approach:

Step 1: We focused on tuning the generic hyperparameters while keeping the specific hyperparameters fixed ($\alpha = 0.1, \beta = 0.1$). We found the optimal generic hyperparameter setting as one that maximized the prediction accuracy on biopsy samples under leave-one-patient-out cross-validation (LOPO CV). This step followed a similar approach commonly employed in other DL algorithms. Some additional details are provided as follows:

- Number of layers: we tried 2, 3, and 4 layers for the shared block and task-specific blocks corresponding to Pro and Inf. 3 layers were the best choice.

- Number of neurons per layer: we tried 64, 128, 256 neurons. The best choice was: the 3 layers in the shared block had 256, 128, and 128 neurons; the 3 layers in task-specific blocks had 128, 128, and 64 neurons.

- Learning rate: we tried 4e-3, 2e-3, 1e-3 for each block. 2e-3 was chosen for the shared block and 1e-3 for the task-specific blocks.

- Batch size: we tried 256, 512, 1024 and chose 1024.

- Number of layers and number of neurons per layer for the auxiliary task corresponding to Neu: to enforce the shared latent representations encoded discriminant features of Neu, we chose a simple one-layer design with 128 neurons.

- Activation function: we adopted the commonly used ReLU function.

- Optimization algorithm: we adopted the commonly used Adam optimizers.

- Number of unlabeled samples: this is a hyperparameter in some semi-supervised learning models. Our model is semi-supervised learning and was designed to include unlabeled samples that survived an uncertainty threshold, PE*. Higher PE* allows for more unlabeled samples to be included, but some included samples have low quality/certainty. Lower PE* results in a smaller number of samples being included. While these samples had relatively higher quality/certainty, the reduced sample size impacts the diversity of the selected samples. To strike a balance, we tuned PE* within a range of 0.3-0.7 and chose PE*=0.44, which resulted in a large number of over 7e+4 unlabeled samples being included. Also, according to the MCMA criterion, this choice projected a high accuracy of over 90%. It is worth noting that our model is not particularly sensitive to this hyperparameter, so that we only performed a rough search than an exhaustive exploration.

Step 2: Under the setting of generic hyperparameters found in step 1, we focused on tuning the specific parameters, which were weights $\alpha$ and $\beta$ in our custom loss function. Two criteria were examined: (1) data-driven criterion which is the prediction accuracy on biopsy samples; (2) knowledge-driven criterion which is the percentage of unlabeled samples whose predictions are in concordance with the two knowledge-based relationships. The same relationships that guided the design of the custom loss function were utilized, namely, Pro and Inf are likely to be low for samples with high Neu; Pro and Inf are likely to be negatively correlated for samples with low Neu.

In detail, criterion (2) was computed as follows: We calculated the percentage of unlabeled samples in $\{i \in neu^-\}$ whose predicted Pro and Inf satisfy one being below 0 and the other above 0. We also calculated the percentage of unlabeled samples in $\{i \in neu^+\}$ whose predicted Pro and Inf are both below 0. The two percentages were averaged to become criterion (2).

Under the two criteria, we tuned $\alpha$ and $\beta$ to maximize criterion (1) while achieving >60% of criterion (2) for each patient under LOPO CV. The chosen threshold of 60% considered that the knowledge-based relationships are qualitative and the approach used to compute criterion (2) was only approximate. Thus, it was deemed reasonable to expect that the majority (>60%) of unlabeled samples would satisfy criterion (2) to align with the knowledge-driven aspect of the evaluation.

Remark: It is important to highlight that, in the tuning of weights in the loss functions of DL algorithms, it is common to rely solely on data-driven criteria. However, our problem presents a unique opportunity to enhance the tuning process by leveraging domain knowledge. This approach proves particularly advantageous as the knowledge can be applied to many unlabeled samples. Given the limited number of biopsy/labeled samples used to compute the data-driven criterion, additionally incorporating the knowledge-driven criterion becomes crucial in helping reduce the risk of overfitting in the tuning results and poor generalization on the test set.

**7. Hyperparameter tuning for competing methods**

The hyperparameter tuning procedure for the competing method was aligned with that employed for BioNet, and can be delineated as follows:

1. **Scale Definition**: Each hyperparameter's scale was appropriately defined to encompass a broad yet relevant range of values.
2. **Random Search**: This step involved exploring random hyperparameter values within the predefined scales, facilitating a comprehensive initial search.
3. **Grid Search**: Subsequent to the random search, a focused grid was established for each hyperparameter. This grid was based on the promising areas identified during the random search, enabling a more detailed exploration.
4. **Default Setting**: In cases where certain hyperparameters exhibited low sensitivity, default values were assigned. This strategy aimed to economize on time and computational resources without compromising the efficacy of the model.
5. **Completion Criterion**: The tuning process was deemed complete when no further performance enhancements were observed on cohort A.
6. **Final Evaluation**: The optimized model was then evaluated on cohort B, a separate test set, to assess its generalization capabilities.

Our competing methods included NN-based methods and classic ML methods. The hyperparameters tuned by the tuning procedure are as follows:

- For NN-based methods including MTL-AAE, AdaMatch, MTL-NN, and NN:

  Learning rate, hidden size, number of hidden layers, batch size, number of shared layers, number of epochs, dropout rate, optimizer, learning rate scheduler.
- For ML methods:
  - SVR: regularization parameter C, kernel, kernel coefficient, independent term in kernel function, degree of the polynomial kernel function.
  - RF: number of trees in the forest, maximum depth of the tree, minimum number of samples required to split an internal node, minimum number of samples required to be at a leaf node, number of features to consider when looking for the best split.

### 8. Computation of the knowledge concordance (KC) metrics overall unlabeled samples

Recall that the domain knowledge indicates two key relationships: (1) Pro and Inf are likely to be negatively correlated for samples with low Neu; (2) Pro and Inf are likely to be low for samples with high Neu. To compute the KC metrics, we first used the trained BioNet_Neu model to stratify unlabeled samples into two groups with low (<0) and high (>0) predicted scores of Neu. Denote these groups by $\{i \in Neu^{-}\}$ and $\{i \in Neu^{+}\}$. Note that these groups included all unlabeled samples within each AOI, not just the unlabeled samples included to train BioNet_ProInf which are samples with high certainty. Furthermore, we computed the KC metric with respect to relationship (1), $KC_{neu^{-}}$, as the percentage of unlabeled samples in $\{i \in Neu^{-}\}$ whose predicted Pro and Inf satisfy one being below 0 and the other above 0. We computed the KC metric with respect to relationship (2), $KC_{neu^{+}}$, as the percentage of unlabeled samples in whose predicted Pro and Inf are both below 0.
